# Identifiability limits of metabolic dynamics from sparse longitudinal metabolomics

**DOI:** 10.64898/2026.08.01.742207

**Authors:** Javad Aminian-Dehkordi, Mohammad Mofrad

## Abstract

Sparse and irregular longitudinal metabolomic measurements can make dynamical quantities numerically estimable without making them scientifically identifiable. We formulate effective-landscape inference as an observation-timescale inverse problem and evaluate its operating characteristics using stochastic systems with known ground truth and longitudinal stool metabolomics from the Integrative Human Microbiome Project. Across simulation ensembles, recovery of a known double-well topology declined sharply as observations became sparse, whereas the tested monostable controls did not generate spurious secondary minima. Findings establish an identifiability hierarchy: global topology requires sufficient within-subject state-space coverage, whereas local return descriptors remain vulnerable to bias from the observation process. The framework therefore shifts the emphasis from assigning stability labels to determining what a sampling design can support. It provides a basis for designing denser longitudinal experiments and for validating clinical and machine-learning studies in which sparse observations might otherwise be overinterpreted as evidence about underlying dynamics.

## 1 Introduction

The gut microbiome is a dynamic ecosystem whose metabolic products respond to diet, inflammation, medication, and other perturbations^1^. Short-chain fatty acids such as butyrate are important host-facing metabolites, but their longitudinal measurements are often sparse relative to microbial and metabolic timescales^2,3^. This mismatch creates an inverse problem: the same observed trajectory may be compatible with multiple latent dynamical systems.

Ecological stability is commonly related to the tendency of a system to return toward an operating state after displacement^4^. A one-dimensional drift field can be represented geometrically through an effective potential, with local minima corresponding to candidate stable states and local curvature describing a restoring tendency^5^. Multiple minima provide an effective representation of alternative stable states^6^, whereas barrier geometry and transition frequency describe global features that require broad state-space coverage^7^. Reliable inference of these global properties generally requires sufficient observations across the relevant state space, including regions separating candidate stable states.

Sparse time series pose two related risks. Global topology may be missed because transitions between basins occur between observations, while local return may be over-interpreted because regressing change on baseline is vulnerable to finite-sample mathematical coupling and measurement error^8^. Accordingly, false-negative and false-positive operating characteristics must be evaluated before landscape or return parameters are assigned biological meaning^9,10^.

We therefore develop a theoretical and computational framework addressing three objectives. First, we quantify how rapidly a known multistable topology becomes undetectable as sampling becomes sparse. Second, we assess when a short-series return coefficient can be distinguished from the negative slope generated under a non-restorative null. Third, we evaluate whether candidate metabolite covariates improve predictive performance after accounting for multiplicity and subject influence. The individual estimators are established; the contribution lies in their integration into a ground-truth operating-characteristic analysis tailored to sparse longitudinal metabolomics.

In this study, the empirical HMP2 trajectories serve as a case study. The resulting analysis separates computable descriptors from biologically identified quantities and translates each inferential limitation into a concrete requirement for future study design. By making the observation process explicit, the framework provides a way to assess whether a planned longitudinal experiment or clinical study contains the temporal information needed to support its intended dynamical claims. More broadly, this work provides a validation layer for aligning sampling schedules, model complexity, and biological interpretation before sparse observations are translated into mechanistic or clinical conclusions.

## 2 Theoretical framework

### 2.1 Stochastic dynamics and effective potential

Let *B*(*t*) *≥* 0 denote measured butyrate intensity and let *z*(*t*) = log[1 + *B*(*t*)] denote the transformed state. At the observation scale, we represent the unresolved high-dimensional metabolic system by the one-dimensional stochastic process^11^

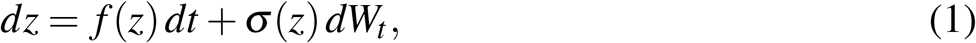

where *f* (*z*) is the conditional drift, *σ* (*z*) summarizes unresolved process variability, and *W*_*t*_ is a Wiener process. This is a coarse-grained statistical representation, not a mechanistic reduction of microbial reactions. In one dimension, a drift potential can be defined as

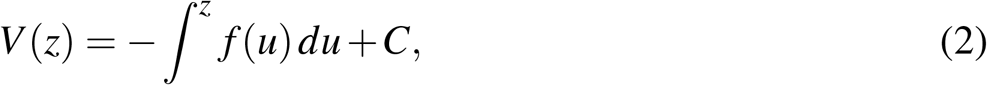

where the arbitrary constant *C* has no dynamical consequence. A point *z*^*\**^ satisfying *f* (*z*^*\**^) = 0 and *f* ^*′*^(*z*^*\**^) *<* 0 corresponds to a local minimum of *V*. Its drift curvature is

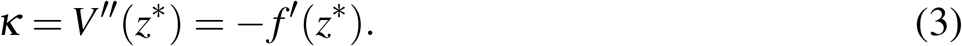

The potential is therefore a geometric representation of the estimated drift. It is not a thermodynamic free-energy landscape, and outside the visited state range it is not constrained by the observations. A direct stationary-density interpretation would additionally require assumptions about diffusion and stationarity that are not invoked here^12,13^.

### 2.2 Observation operator and discrete return model

Observations occur at irregular times *t*_*i*_ and may include measurement error,

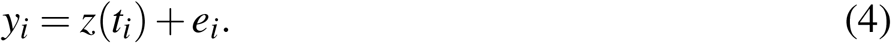

For sufficiently short intervals, *E*[Δ*z*_*i*_ | *z*_*i*_] *≈ f* (*z*_*i*_)Δ*t*_*i*_, but this approximation weakens when the sampling interval spans relaxation or switching timescales^14^. The low-parameter stepwise model

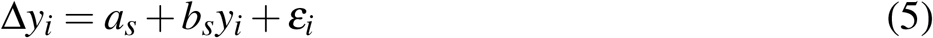

therefore describes transitions per observation step. If it were a valid local linearization with comparable intervals and negligible observation error, 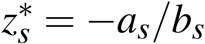 and *−b*_*s*_ would describe an apparent local restoring strength.

Measurement error makes the limitation explicit. Because

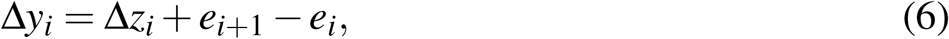

the same error *e*_*i*_ enters the predictor positively and the response negatively. Under independent errors, this contributes *−*Var(*e*_*i*_) to Cov(*y*_*i*_, Δ*y*_*i*_) and biases the fitted slope downward. Short random walks also yield predominantly negative finite-sample slopes even when observation error is absent. Consequently, a negative *b*_*s*_ must be benchmarked against appropriate null processes before it is interpreted as recovery or resilience.

### 2.3 Topology and identifiability

Multistability is treated as a testable property of the reconstructed observed-range potential. Evidence requires at least two separated minima, a non-negligible intervening barrier, and reproducibility under perturbation of the observed transition set. Failure to meet these criteria is a non-detection at the observation scale; it does not exclude faster switching, unvisited basins, or multivariate mechanisms hidden by the one-dimensional projection.

This distinction motivates two complementary validation layers. Simulations with known topology quantify sensitivity and specificity across sampling and noise regimes. Random-walk and measurement-error nulls quantify whether an apparent local return coefficient is distinguishable from a non-restorative process. Figure 1 summarizes the point at which statistical validation constrains biological interpretation.

**Figure 1:**
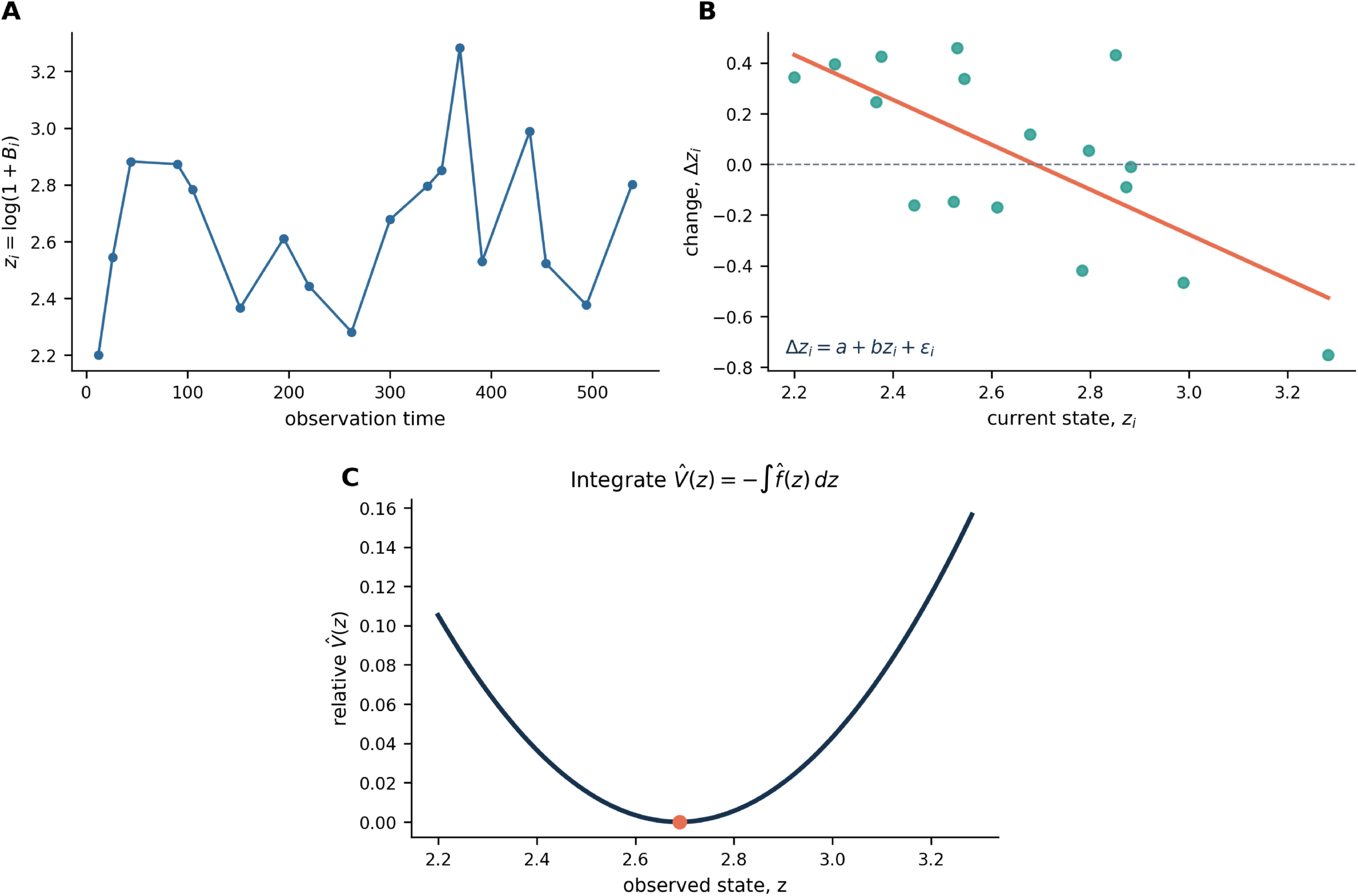
Effective-landscape inference at the observation timescale. An illustrative, independently generated trajectory shows the analysis. (A) Sparse transformed observations define the available state range. (B) Consecutive changes are related to the current state to estimate transition-scale drift. (C) Integrating the estimated drift gives a relative effective potential only over the visited range.

## 3 Results

### 3.1 Sparse sampling obscures known bistability

We first simulated a bistable double-well system and a monostable Ornstein–Uhlenbeck (OU) control. A selected trajectory reproduced the expected illustration: the double-well system yielded two inferred minima under dense sampling and one under sparse sampling, whereas the OU process remained single-well (Figure 2). Because a single trajectory can be unrepresentative, inference was then repeated for 200 trajectories at every combination of five noise amplitudes and five sampling strides.

**Figure 2:**
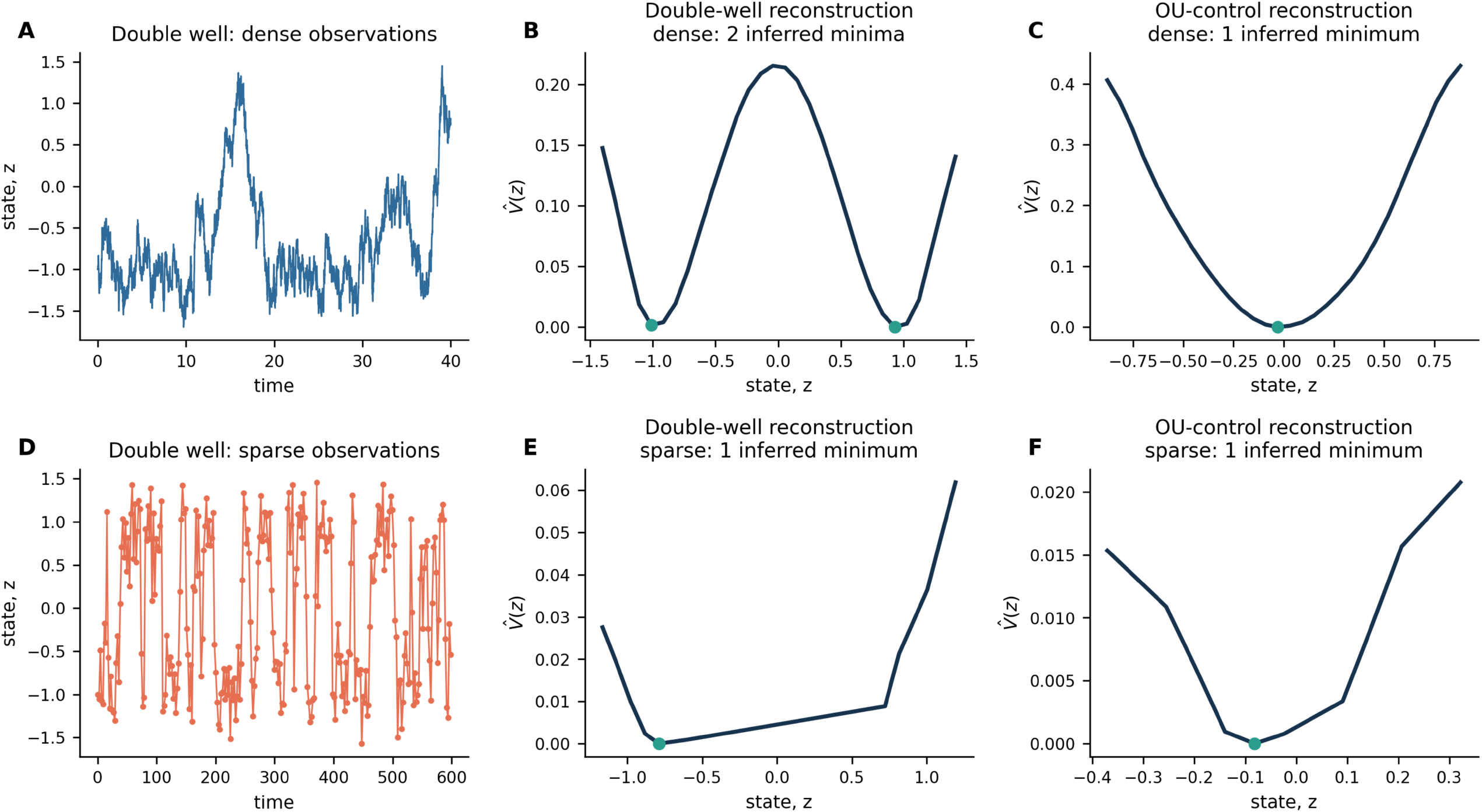
Selected-trajectory demonstration of sampling-induced topology loss. Dense and sparse observations of the same stochastic systems were reconstructed using binned drift and numerical integration. The selected double-well trajectory yields two inferred minima when dense and one when sparse; the OU control remains single-well. Ensemble operating characteristics are reported separately in Figure 3.

At *σ* = 0.6, the two-minimum topology was recovered in 200/200 double-well trajectories at stride 50, 191/200 at stride 100, 89/200 at stride 150, 1/200 at stride 225, and 0/200 at stride 300 (Figure 3). The corresponding 95% Wilson interval at stride 225 was 0.09–2.78%. Across the full control grid, no OU trajectory produced a spurious second reconstructed minimum (0/5,000; pooled 95% Wilson upper bound 0.08%). Thus, under this reconstruction procedure, the known double-well topology becomes practically unrecoverable once observation intervals exceed the simulated switching and relaxation scales. The OU result supports specificity within the tested monostable family (not universal specificity against all monostable processes).

**Figure 3:**
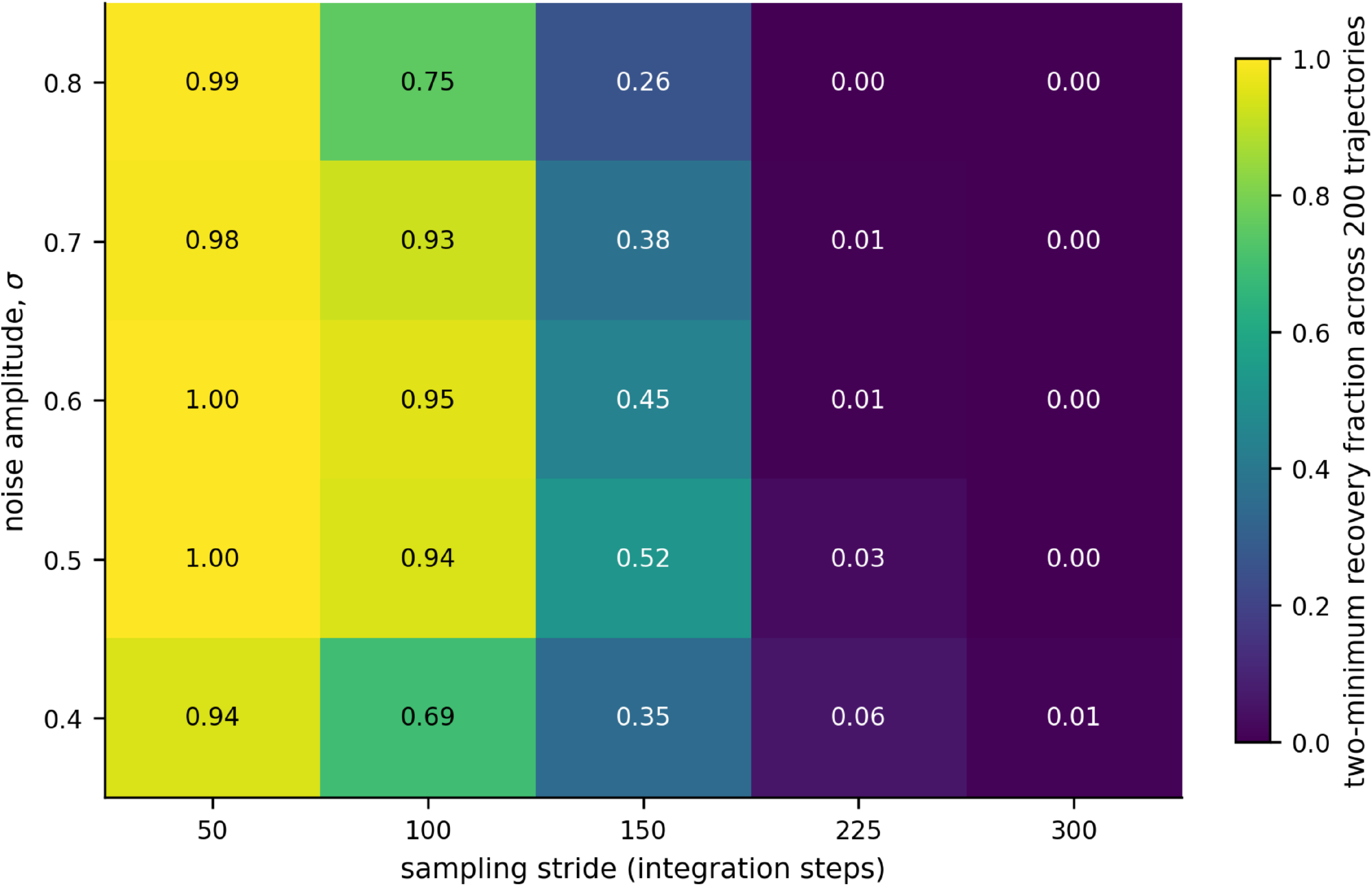
Recovery of known multistability across noise and sampling regimes. Each cell reports the fraction of 200 independently simulated double-well trajectories in which at least two reconstructed minima were detected. The separately evaluated OU control produced no spurious second minimum in any of 5000 trajectories; this negative-control result and its Wilson bounds are reported in the Supplementary Information.

### 3.2 HMP2 cohort and sampling intervals

The canonical data were reconstructed from the HMP2 sample metadata/SCFA table and metabolomics intensity matrix^15^. The source had 546 samples from 106 subjects. Ten subjects had at least six usable consecutive butyrate transitions; four had at least seven and formed the primary cohort. Each retained subject had eight observations and seven transitions. The retained observation gaps were heterogeneous: 14–124 days overall, with a pooled median of 41 days. Subject-level medians ranged from 28 to 42 days (Supplementary Table S1 and S2). These intervals do not support direct cross-subject ranking of step-based return coefficients. An irregular-time OU sensitivity fit is therefore reported only as an exploratory check in the Supplementary Table S6.

### 3.3 Multistability detection across subjects

Nonparametric drift reconstruction produced no full-data topology satisfying the primary separation and barrier criteria. Bootstrap multi-well reproducibility was zero for every subject, and no subject was classified as multistable (Figure S2). This result was unchanged across all 27 combinations of separation, barrier, and reproducibility thresholds evaluated in sensitivity (Supplementary S5). The simulation results show that a true double-well process can yield the same observed classification when sampled sparsely.

### 3.4 Apparent return coefficients

The fitted stepwise coefficients were *−*1.356, 0.536, *−*1.034, and *−*0.794 for C3032, C3037, H4001, and H4004 subjects, respectively. Thus, three of four point estimates were negative. All paired-transition bootstrap intervals crossed zero (Figure 4), and the bootstrap is interpreted as a resampling-stability diagnostic because seven transitions cannot support a reliable dependence-aware block bootstrap.

**Figure 4:**
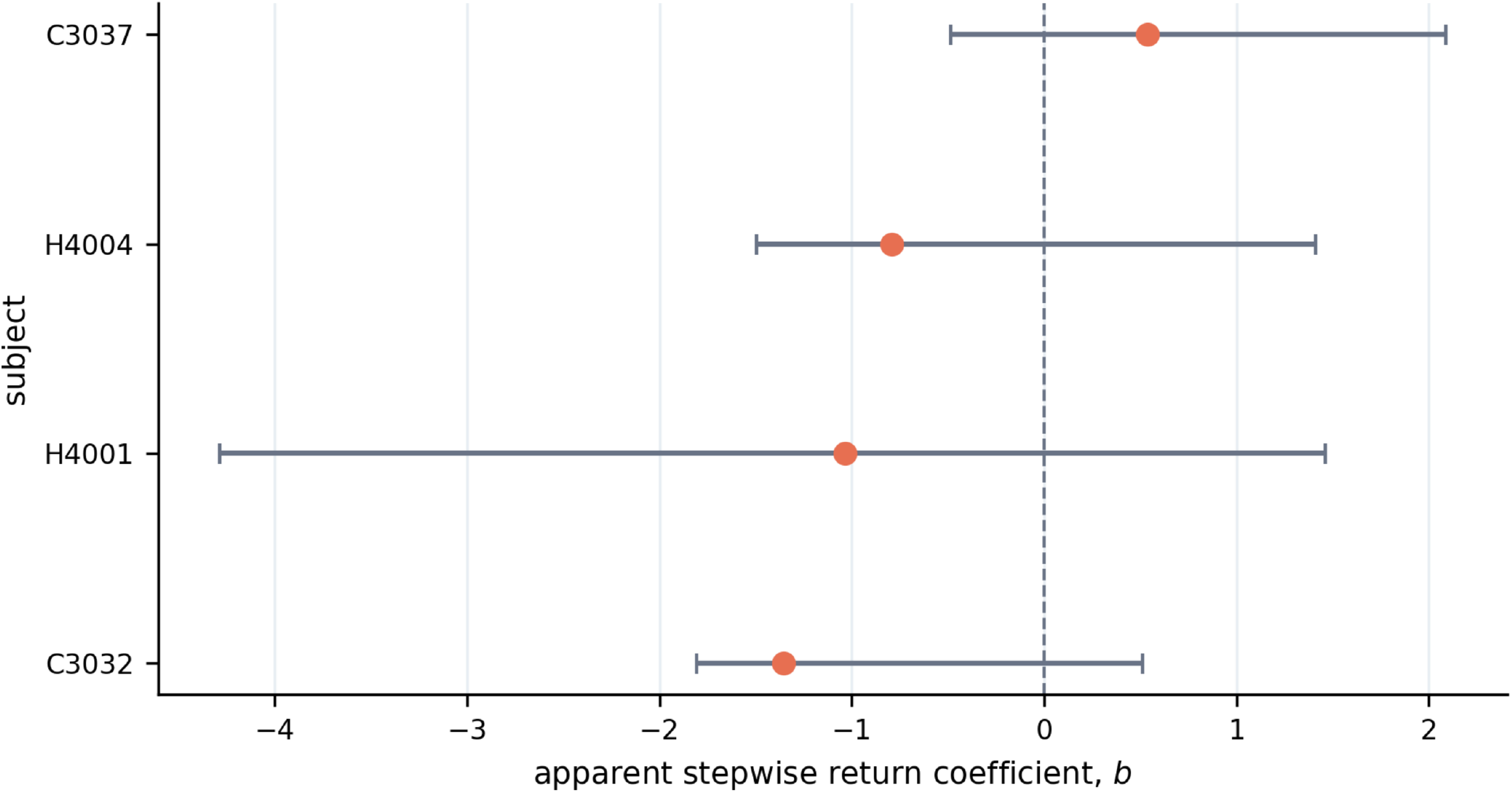
Apparent stepwise return coefficients. Points show subject-specific estimates from Δ*z*_*t*_ = *a*_*s*_ + *b*_*s*_*z*_*t*_ + *ε*_*t*_. Horizontal ranges are raw paired-transition bootstrap percentiles and are descriptive. Every range crosses zero.

More importantly, negative slopes were common under a random-walk null. Among 50,000 simulated eight-observation random walks, 92.6% produced *b <* 0 without observation error. This increased to 97.8% when measurement-error standard deviation equalled innovation standard deviation. The median null slope changed from *−*0.502 to *−*0.806 across these settings (Figure 5 and Supplementary S3). Only C3032 was in the lower 5% tail of the no-measurement-error null, and it was no longer below that threshold under moderate measurement error. Consequently, the coefficients are reported as apparent stepwise return coefficients, not estimates of subject resilience or landscape curvature.

**Figure 5:**
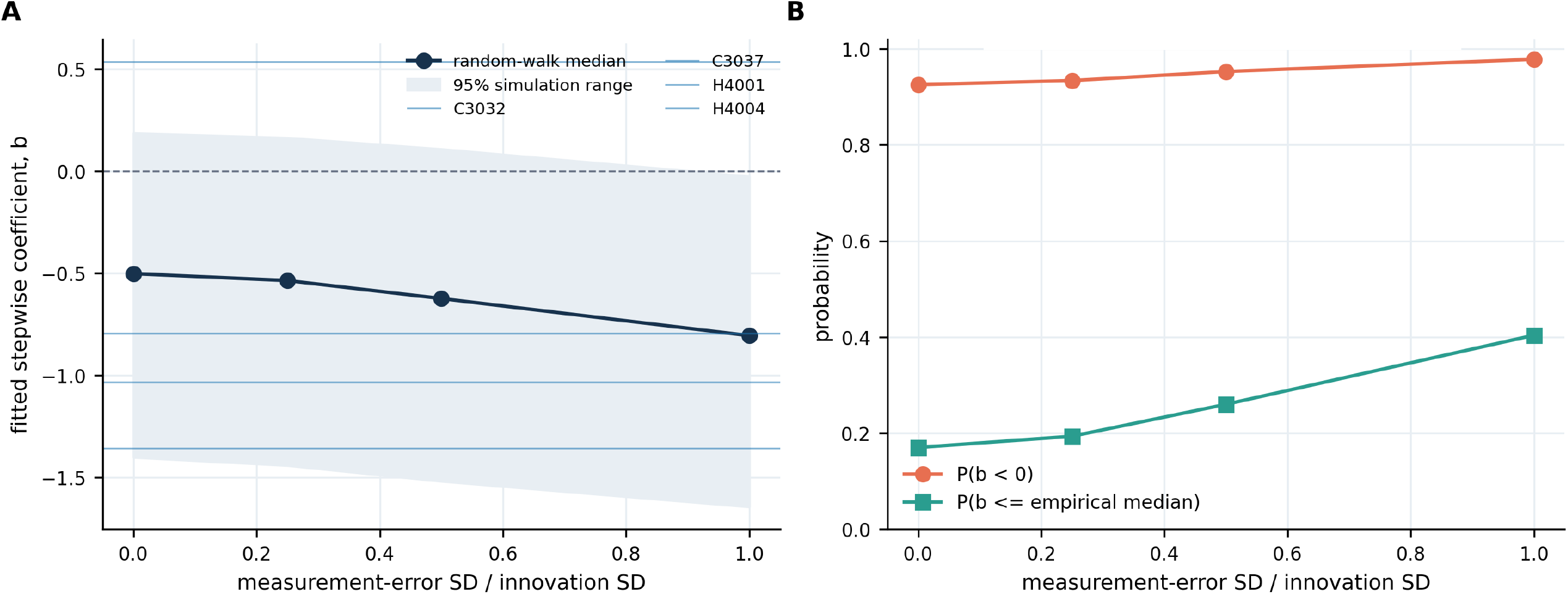
Random-walk and measurement-error operating characteristics. Short-series regression of Δ*z*_*t*_ on *z*_*t*_ frequently produces a negative coefficient in the absence of restorative dynamics. Observation error further shifts the null distribution in the negative direction.

**Figure 6:**
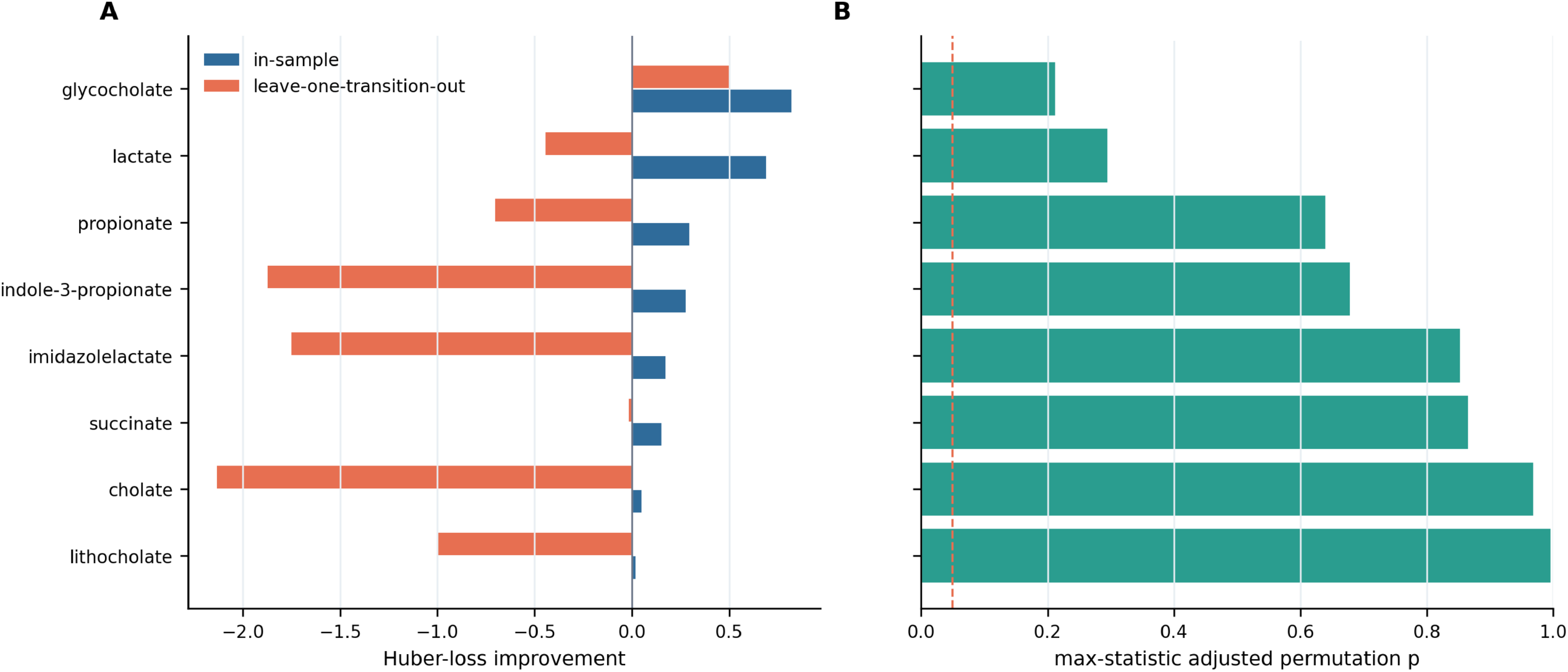
Robustness of exploratory metabolite associations. In-sample improvement, leave-one-transition-out improvement, and family-wise permutation probabilities are shown for all eight candidates. No candidate met the multiplicity-adjusted threshold.

We also compared the prespecified linear drift with a subject-specific quadratic alternative. Although the additional term necessarily reduced in-sample residual error, small-sample corrected AIC favored the linear model in all four subjects, and leave-one-transition-out error favored the linear model in three of four. For C3037, the quadratic predictive improvement was negligible (RMSE 0.568 versus 0.576). This diagnostic provides no stable evidence that a quadratic drift is supported by seven transitions, but the low power also means that linearity should be treated as a working approximation rather than an established biological law (Supplementary Table S4).

### 3.5 Metabolite associations remain hypothesis-generating

Eight candidate metabolites were screened using a subject-fixed-effects ARX model with subject-specific intercepts and state slopes. Glycocholate and lactate produced the largest in-sample Huber-loss improvements. Glycocholate retained a positive leave-one-transition-out loss improvement, whereas lactate did not. Neither survived the exact circular-shift maximum-statistic correction across eight candidates (adjusted *P* = 0.212 and 0.294, respectively). Omitting H4001 reversed both coefficients and eliminated their loss improvements. The same two metabolites ranked first in the broader 10-subject, six-transition sensitivity cohort, but this concordance does not overcome the lack of adjusted evidence in the primary cohort. Accordingly, the screen does not identify metabolite modulators; it provides candidate associations for future targeted studies (Supplementary S7).

## 4 Discussion

Longitudinal metabolomics can potentially reveal if a metabolic variable returns toward a characteristic level after fluctuation, shifts between alternative persistent states, or changes without a detectable restoring tendency. The effective-landscape framework summarizes these possibilities geometrically: a potential minimum represents a concentration range toward which the system tends to return, curvature describes the local strength of that tendency, and multiple separated minima may indicate alternative dynamical states. Our results show that the interpretability of these features depends critically on the sampling process.

Global landscape structure, such as the presence of multiple wells, requires observations covering a sufficiently broad range of the system and capturing movement between dynamical regions. Local return descriptors require less coverage, but can still be biased by short time series, measurement error, irregular sampling intervals, and the mathematical coupling created when change is regressed on its baseline value. We therefore distinguish three categories: quantities whose operating characteristics can be evaluated using ground-truth simulations, quantities that are computable but only descriptively interpretable, and quantities that remain unsupported under the present design (Table 1). This distinction is more informative than assigning a single label of stable or unstable to a sparsely observed biological system.

**Table 1:**
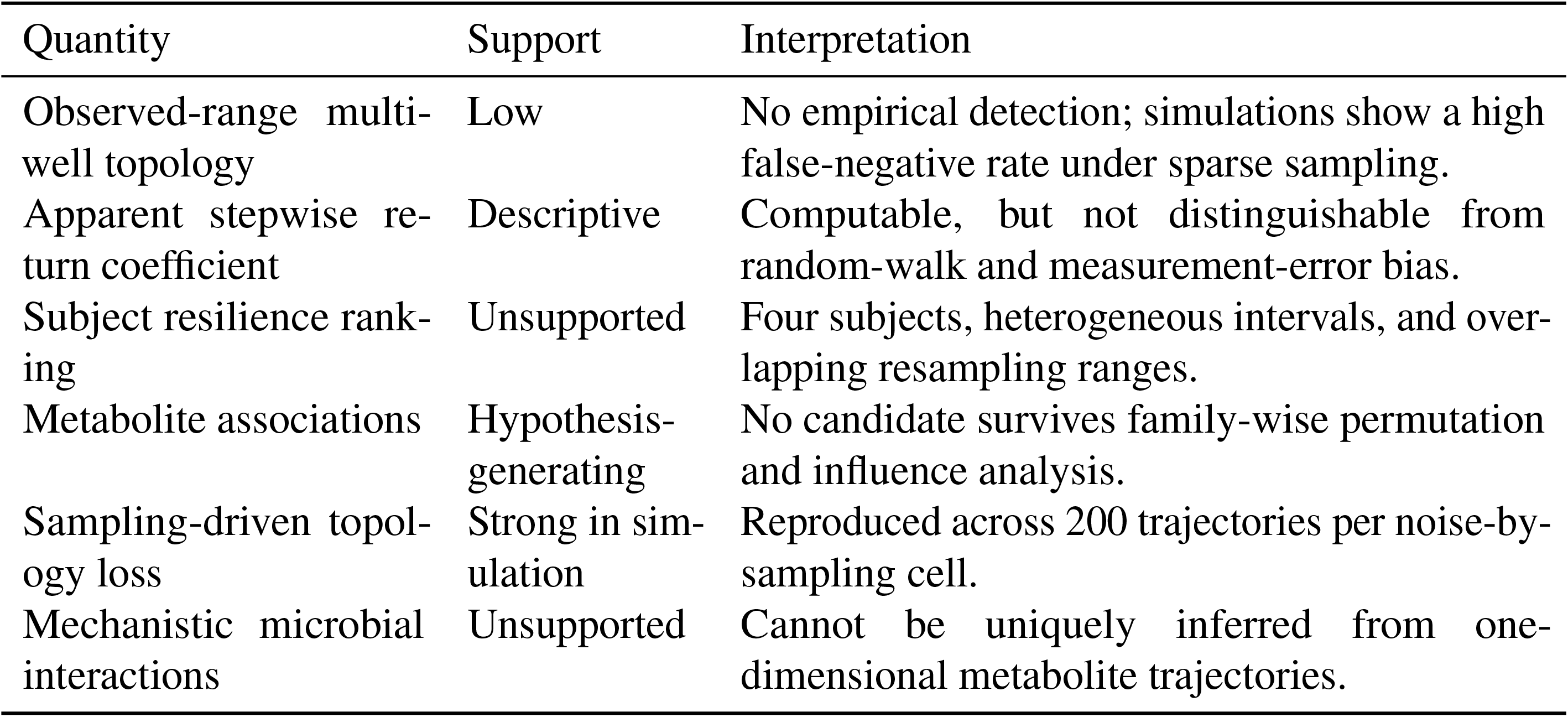
Interpretation of dynamical quantities under the present sampling design.

The ground-truth simulations show the limitation affecting global landscape structure (Figure 3). Sparse sampling progressively obscured the two known states of the double-well system, despite no change in its underlying dynamics. Conversely, the tested OU controls, which represent a single restoring state, did not produce spurious secondary minima. These models therefore assess complementary errors: failure to detect genuine multistability and false detection under a specified monostable null. Neither is intended as a mechanistic model of gut metabolism, and the OU results do not establish specificity against every nonlinear monostable process. Although the empirical variability and observation intervals informed the reference simulation setting, recovery was evaluated across a broader grid of noise amplitudes and sampling intervals. The observed loss of topology therefore does not depend on selecting a single HMP2-matched parameter combination. The supported conclusion is that sparse observation can conceal genuine multi-well structure, while the reconstruction did not manufacture such structure in the monostable controls examined here.

The local return analysis exposes a different identifiability problem. A negative regression of change on baseline is common in an eight-observation random walk, and observation error strengthens the negative dependence because the baseline error appears with the opposite sign in the subsequent change. Thus, even though *a*_*s*_, *b*_*s*_, and the algebraic equilibrium *−a*_*s*_*/b*_*s*_ can be fitted, their existence as numerical outputs does not establish a biological equilibrium or recovery rate. The empirical slopes cannot support subject-specific resilience claims or rankings. This result refines the theoretical landscape framework: local curvature remains a valid property of a known drift field, but the present observations do not identify that curvature separately from finite-sample and measurement processes.

The fitted coefficients quantify change per observed transition, not recovery per unit time. Because observation gaps ranged from 14 to 124 days, the step-based coefficients cannot be compared directly as physical recovery rates across subjects. Time-normalized regressions and irregular-time OU fits were examined as sensitivity analyses, but remained weakly identified with only seven transitions per subject and included a boundary estimate. Likewise, paired-transition bootstrap ranges describe sensitivity to the observed transitions.

The empirical landscape reconstructions should be interpreted under the same logic. No subject showed a reproducible observed-range multi-well topology under the primary criteria or any of the 27 threshold combinations. This constrains what is detected at the sampling timescale, but it does not demonstrate global monostability.

Also, the metabolite extension shows how the framework can be expanded without converting association into mechanism. Subject-specific intercepts and state slopes, together with within-subject centering, separate the common metabolite coefficient from simple between-subject mean differences. Nevertheless, only four trajectories inform that coefficient. Glycocholate and lactate improved in-sample loss, but neither survived family-wise correction and both were sensitive to omission of H4001; lactate also failed to improve leave-one-transition-out prediction. These features should therefore be viewed as hypotheses for targeted perturbation studies, not as established modulators.

The theoretical framework points directly to a stronger experimental design. Resilience studies should sample more frequently than the expected relaxation and switching timescales, obtain technical replicates or an external estimate of observation error, record perturbation onset and resolution, and follow enough independent subjects to estimate population heterogeneity. In the simulated double-well system at *σ* = 0.6, recovery decreased from 95.5% with 601 observations at an effective interval of 1.0 model-time unit to 44.5% with 401 observations at an interval of 1.5 and 0.5% with 267 observations at an interval of 2.25. If the per-subject detection probability under a specified design is *p* and the population prevalence is *q*, the probability of at least one detection among *N* independent subjects is 1*−*(1*− pq*)^*N*^; this separates within-subject detection from estimation of population prevalence. Controlled interventions would help distinguish restoration from regression to the mean, while paired metagenomic, metabolomic, and flux measurements could test whether the one-dimensional drift reflects identifiable community mechanisms.

These considerations extend beyond effective-landscape reconstruction. In experimental and clinical studies, the target dynamical quantity should be specified before data collection, and the observation schedule should be evaluated against the timescale on which that quantity can change. More generally, imperfect observation can propagate from uncertainty in state estimation to loss of decision quality by restricting the set of interventions that can be justified from the available data, as formalized in a complementary feasible-set framework for microbiome intervention under observation loss^16^. Increasing the number of participants cannot compensate for insufficient within-subject resolution when the objective is to identify recovery, switching, or transient responses to perturbation.

The same distinction is relevant to machine-learning analyses of longitudinal biological data. Flexible models may extract predictive associations from sparse and irregular measurements, but predictive performance alone does not establish that the underlying transition dynamics are identifiable; apparent signal may still reflect between-subject structure, measurement coupling, interpolation across long intervals, or other features of the observation process. Longitudinal machine-learning studies would therefore benefit from sampling-density simulations, subject-level validation, and sensitivity analyses for irregular timing and observation error. In this sense, the present framework can serve as a design and validation layer for determining whether a dataset contains the temporal information required for the biological or clinical interpretation assigned to a fitted model.

In summary, the value of effective-landscape inference in sparse metabolomics lies not in forcing a state assignment, but in formalizing what an observation schedule permits. The present analysis provides strong simulation evidence for sampling-driven topology loss, a rigorous warning against interpreting short-series return slopes as resilience, and a reproducible template for evaluating future data. Biological resilience, subject rankings, and metabolite modulation remain questions for denser and prospectively designed studies.

## 5 Methods

### 5.1 Data source and cohort selection

We used publicly available longitudinal stool metabolomics from HMP2^15^. The analysis table was constructed from a 546-sample metadata/SCFA table and a metabolomics intensity matrix by a one-to-one join on sample identifier.

Samples with a valid receipt date and non-negative butyrate intensity were ordered within subject. A usable transition connected two consecutive observations. The primary rule required at least seven transitions, yielding C3032, C3037, H4001, and H4004. The configured identifiers are checked programmatically against the rule-derived set. A six-transition threshold yielding 10 subjects was used for a secondary metabolite-ranking sensitivity analysis.

Butyrate intensity *B*_*t*_ was transformed as *z*_*t*_ = log(1 + *B*_*t*_). The parametric return analysis used unsmoothed observations. Smoothing was applied only within the explicitly defined nonparametric landscape reconstruction.

### 5.2 Stepwise return model

For consecutive observations,

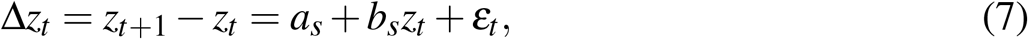

where *a*_*s*_ and *b*_*s*_ are subject-specific. If the coefficient represented a valid local linearization of a restorative process, *−b*_*s*_ would correspond to a per-observation-step curvature. Because observation intervals differ and the random-walk null produces substantial negative bias, we treat *b*_*s*_ as a descriptive apparent return coefficient.

Transitions were resampled with replacement 1,000 times within subject. Percentile ranges are reported as paired-transition resampling diagnostics. They are not interpreted as nominal time-series confidence intervals^17^.

As a functional-form diagnostic, the subject-specific linear model was compared with 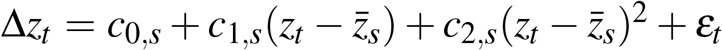. We report in-sample RMSE, *R*^2^, small-sample corrected AIC, and leave-one-transition-out RMSE for both specifications. Because each subject has only seven transitions, the quadratic comparison is used to detect gross lack of fit and is not interpreted as selection of a mechanistic nonlinear drift.

### 5.3 Random-walk and measurement-error null

We simulated 50,000 independent random-walk trajectories with eight observations,

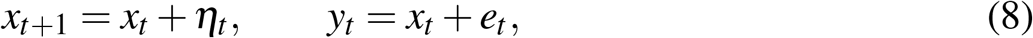

where *η*_*t*_ ~ *N* (0, 1) and *e*_*t*_ ~ *N* (0, *r*^2^). The measurement-error ratios were *r* ∈ {0, 0.25, 0.5, 1}. The empirical regression was refitted to each simulated observed trajectory. We report the null median, 95% simulation range, probability of a negative slope, and subject-specific lower-tail probabilities. This analysis addresses small-sample mathematical coupling and observation-error bias^10,18^.

### 5.4 Nonparametric landscape and multistability criteria

Transitions were binned into five equal-width bins over the subject’s observed *z*_*t*_ range. Bins with at least one transition were retained. Mean drift was interpolated to 241 grid points, smoothed with a 17-point moving-average window, integrated using the trapezoidal rule, and the resulting potential was smoothed with an 11-point window. A candidate pair of minima was required to be separated by at least 20% of the observed state range and by a barrier at least 5% of the potential range. A subject was classified as multistable only if the full reconstruction qualified, at least 80% of 1,000 bootstrap reconstructions were valid, and at least 80% of valid reconstructions were multi-well. Sensitivity analysis crossed state-separation fractions {0.15, 0.20, 0.25}, barrier fractions {0.025, 0.05, 0.1}, and reproducibility fractions {0.7, 0.8, 0.9}, using 500 resamples per setting.

### 5.5 Stochastic simulation operating characteristics

The double-well model used

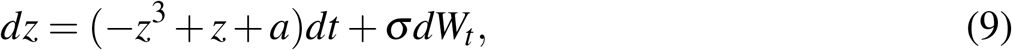

with *a* = 0, and the monostable control used

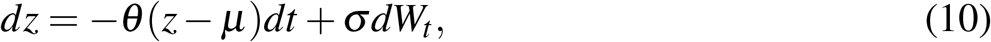

with *θ* = 1.2 and *µ* = 0. Euler–Maruyama integration used *dt* = 0.01 and total duration 600^12,14^. Ensemble sensitivity crossed *σ ∈ {*0.4, 0.5, 0.6, 0.7, 0.8} with sampling strides {50, 100, 150, 225, 300} and 200 trajectories per cell. Detection fractions are accompanied by Wilson intervals.

For this ground-truth topology experiment, recovery was defined structurally as at least two interior local minima in the reconstructed, smoothed potential. This isolates whether the known two-well geometry remains visible after subsampling.

### 5.6 Metabolite association screen

For each of eight exact-name metabolites, we fitted

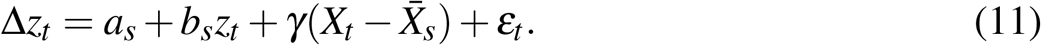

The model includes subject-specific intercepts and slopes and a common within-subject metabolite coefficient. Models were fitted by least squares and ranked by improvement in Huber loss relative to the subject-specific baseline^19^. Robustness was evaluated using leave-one-transition-out prediction, leave-one-subject-out influence analysis, and exact within-subject circular shifts. All 7^4^ = 2401 shift combinations were evaluated for the primary cohort. The maximum loss improvement across all eight candidates in each permutation provided family-wise adjusted probabilities.

### 5.7 Elapsed-time and exposure sensitivity analyses

Subject-level sampling intervals are reported directly. As a sensitivity analysis, we fitted both Δ*z*_*t*_*/*Δ*t*_*t*_ = *α*_*s*_ + *β*_*s*_*z*_*t*_ + *ε*_*t*_ and the irregular-time OU conditional mean

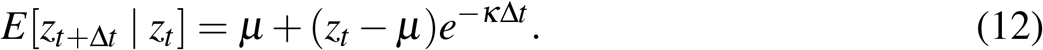

These fits are exploratory because each subject contributes seven transitions. Antibiotic, diarrhea, and hospitalization variables were parsed as categorical fields. For antibiotics, we repeated the stepwise fit after excluding transitions for which either endpoint was antibiotic-positive.

## Data availability

The source metabolomics and metadata are available through HMP2/IBDMDB. The submitted reproducibility package contains the source tables used here, a reconstructed analysis table, and all derived result tables required to regenerate the manuscript.

## Code availability

The code and scripts used to reproduce the numerical analyses and figures will be shared at https://github.com/mofradlab/midloss.

## Competing interests

The authors declare no competing interests.

## Notes

https://github.com/mofradlab/resilience-inference

### Competing Interest Statement

The authors have declared no competing interest.

### Summary of Updates

This manuscript has been substantially revised to strengthen its theoretical framework, statistical validation, and interpretation of dynamical inference from sparse longitudinal metabolomic data. Major revisions include expanded stochastic simulations examining the effects of sampling frequency and noise on the detection of multistability, including systematic sensitivity analyses across multiple parameter combinations. We introduced random-walk and measurement-error null models to evaluate potential biases in estimating local recovery dynamics. These analyses demonstrate that apparent restoring tendencies can arise from short time series even in the absence of genuine restorative dynamics. We also reconstructed and verified the HMP2 dataset, clarified subject selection criteria and sampling intervals, and added sensitivity analyses comparing alternative time-dependent models. The metabolite association analysis has been strengthened through cross-validation, subject-level influence assessments, and multiple-comparison corrections. The manuscript's conclusions have been revised accordingly. Rather than claiming that metabolic resilience can be reliably quantified from the available human data, the study now emphasizes the identifiability limits imposed by sparse longitudinal sampling. The revised framework distinguishes between dynamical properties that can be estimated numerically and those that can be supported scientifically, providing guidance for the design and interpretation of future longitudinal studies.

