## Supplementary Information for "Identifiability limits of metabolic dynamics from sparse longitudinal metabolomics"

### S1 Data sources, cohort construction, and sampling intervals

The empirical input was constructed from two HMP2-derived source files[1]. Eight metabolites were selected: cholate, glycocholate, imidazolelactate, indole-3-propionate, lactate, lithocholate, propionate, and succinate. The joined table contained 546 unique samples from 106 subjects. The primary analysis required at least seven usable consecutive butyrate transitions per subject.

**Table S1:** Primary empirical cohort and observation intervals.

| Subject | Samples | Transitions | Median gap (days) | Gap range (days) |
|---|---|---|---|---|
| C3032 | 8 | 7 | 28 | 14–124 |
| C3037 | 8 | 7 | 41 | 26–85 |
| H4001 | 8 | 7 | 42 | 14–85 |
| H4004 | 8 | 7 | 39 | 16–84 |

The resulting cohort flow was:

**Table S2:** Cohort flow.

| Stage | Subjects |
|---|---|
| Joined HMP2 source | 106 |
| At least six usable transitions | 10 |
| At least seven usable transitions | 4 |
| Primary analysis cohort | 4 |

### S2 Subject-level sampling and return estimates

The paired transition bootstrap is not presented as a dependence-aware confidence interval. With seven transitions, a block bootstrap would contain too few effective blocks for stable uncertainty estimation. We retain paired resampling only to show how unstable estimates are to the observed transition set.

As a goodness-of-fit diagnostic, the linear return model was compared with a quadratic drift after centering the state within subject. The extra term reduced in-sample RMSE in every subject, as expected for a nested model, but small-sample corrected AIC favored the linear model in all four subjects. Leave-one-transition-out RMSE favored the linear model in

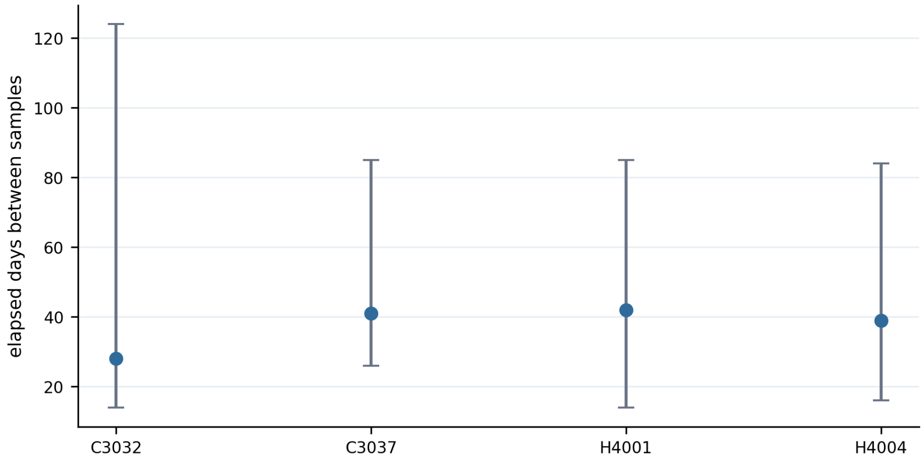


**Figure S1: Subject-specific sampling intervals.** Points show medians; bars span minimum to maximum elapsed days.

**Table S3:** Stepwise return estimates and raw paired-bootstrap percentile ranges. No interval excludes zero.

| Subject | $b$ | 2.5% | 97.5% | $-b$ | Valid resamples |
|---|---|---|---|---|---|
| C3032 | -1.356 | -1.804 | 0.511 | 1.356 | 1000/1000 |
| C3037 | 0.536 | -0.487 | 2.087 | -0.536 | 1000/1000 |
| H4001 | -1.034 | -4.288 | 1.460 | 1.034 | 1000/1000 |
| H4004 | -0.794 | -1.495 | 1.409 | 0.794 | 1000/1000 |

three subjects. The small quadratic advantage for C3037 was 0.008 RMSE units and was not accompanied by AICc support.

**Table S4:** Linear-versus-quadratic return-model diagnostic. Lower AICc and leave-one-transition-out (LOTO) RMSE indicate better support.

| Subject | Linear AICc | Quadratic AICc | Linear LOTO RMSE | Quadratic LOTO RMSE |
|---|---|---|---|---|
| C3032 | -11.79 | -4.97 | 0.325 | 0.585 |
| C3037 | -7.38 | -2.33 | 0.576 | 0.568 |
| H4001 | 1.69 | 7.58 | 1.022 | 7.858 |
| H4004 | 7.22 | 14.07 | 1.215 | 6.495 |

With seven transitions, higher-order drift is weakly identified and quadratic leave-one-out predictions can be dominated by a single high-leverage state. The linear specification is retained as the lower-variance descriptive approximation.

### S3 Random-walk and measurement-error null

For each measurement-error ratio, 50,000 eight-observation random walks were simulated and fitted using the empirical regression. The innovation standard deviation was fixed at one; the measurement-error standard deviation was varied relative to this innovation scale.

**Table S5:** Random-walk null distribution of the fitted stepwise coefficient.

| $\sigma_e/\sigma_\eta$ | Median $b$ | 95% range | $P(b < 0)$ | $P(b \leq -0.914)$ |
|---|---|---|---|---|
| 0.00 | -0.502 | [-1.401, 0.186] | 0.926 | 0.170 |
| 0.25 | -0.536 | [-1.441, 0.163] | 0.934 | 0.193 |
| 0.50 | -0.624 | [-1.519, 0.108] | 0.953 | 0.260 |
| 1.00 | -0.806 | [-1.644, -0.026] | 0.978 | 0.404 |

These simulations show that the sign and magnitude of the observed coefficients are compatible with a non-restorative process over a broad error range. Technical replicates or external error calibration would be needed to separate process and observation noise.

### S4 Irregular-time sensitivity

**Table S6:** Exploratory elapsed-time fits. Boundary estimates and sign changes illustrate weak identification.

| Subject | Daily linear slope | OU $\kappa$/day | OU SSE | Note |
|---|---|---|---|---|
| C3032 | -0.0288 | 0.0990 | 0.529 | Positive return rate |
| C3037 | 0.0200 | -0.0038 | 1.193 | Non-restorative point estimate |
| H4001 | -0.0254 | 0.1000 | 3.289 | Upper-bound estimate |
| H4004 | -0.0118 | 0.0281 | 6.329 | Positive return rate |

The fitted OU rate for H4001 reached the prespecified upper bound. With seven transitions, these models cannot support precise physical-time comparisons and are reported only to show sensitivity to interval treatment.

### S5 Multistability criteria and threshold sensitivity

The primary topology settings were:

- five equal-width state bins and minimum bin count of one;
- 241 interpolation-grid points;
- drift smoothing window of 17 grid points;
- potential smoothing window of 11 grid points;
- minimum well separation of 0.20 of the observed state range;
- minimum barrier of 0.05 of the reconstructed potential range;
- minimum multi-well reproducibility of 0.80;
- 1,000 primary bootstrap replicates and minimum valid fraction of 0.80.

The threshold sensitivity crossed three separations, three barriers, and three reproducibility requirements, yielding 27 settings per subject. Five hundred bootstrap replicates were used for each setting. No subject was classified as robustly multistable under any combination.

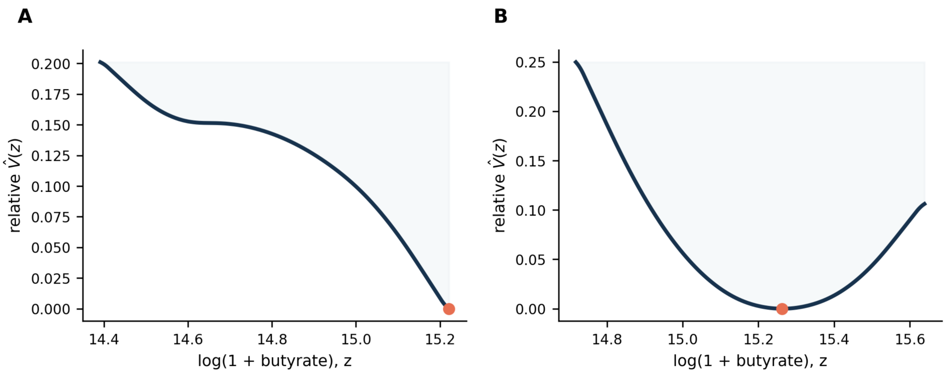


**Figure S2: Observed-range empirical landscape reconstructions.** For each subject, consecutive transitions were binned to estimate drift, smoothed, and numerically integrated to obtain a relative effective potential. Panels show representative (A) boundary-supported and (B) interior-minimum reconstructions. No subject met the primary multi-well criteria.

### S6 Toy-model calibration and ensemble sensitivity

The illustrative simulation used $dt = 0.01$, duration 600, $\sigma = 0.6$, double-well tilt $a = 0$, OU rate $\theta = 1.2$, and sparse stride 225. These values define a study-relevant stress test. The earlier calibration based on the coefficient of variation of signed increments was removed because the increment mean can be near zero, making that ratio unstable.

The ensemble analysis used 200 independent trajectories per cell. For this ground-truth experiment, two-minimum recovery was defined as at least two interior local minima in the reconstructed potential. At $\sigma = 0.6$, double-well recovery fractions were 1, 0.955, 0.445, 0.005, and 0 for strides 50, 100, 150, 225, and 300, respectively. Across the full $5 \times 5$ grid, double-well recovery ranged from 0 to 1 and declined strongly with sampling stride.

**Table S7:** Sampling-density operating characteristics for the double-well model at $\sigma = 0.6$. Observation counts include the initial state and are specific to the simulated duration of 600 model-time units.

| Sampling stride | Effective interval | Observations | Two-minimum recovery |
|---:|---:|---:|---:|
| 50 | 0.50 | 1201 | 1 |
| 100 | 1.00 | 601 | 0.955 |
| 150 | 1.50 | 401 | 0.445 |
| 225 | 2.25 | 267 | 0.005 |
| 300 | 3.00 | 201 | 0 |

The number of subjects and the number of observations per subject answer different design questions. Increasing subjects improves precision for population prevalence or heterogeneity, but it does not recover unobserved within-subject switching. If $p$ is the per-subject detection probability under a prespecified schedule and $q$ is the prevalence of the topology, the probability of at least one detection in $N$ independent subjects is $1-(1-pq)^N$. Translation to a human-study sample size therefore requires a pilot-based estimate of the relevant relaxation time, switching time, noise, and prevalence; the simulation counts above should not be transferred directly to HMP2.

No OU control trajectory produced a spurious second minimum in any cell (0/200 per cell; 0/5,000 pooled). For a single cell, the 95% Wilson interval for 0/200 is 0–1.88%; the pooled 95% Wilson interval is 0–0.077%.

### S7 ARX robustness and multiple testing

**Table S8:** Primary metabolite screen. Positive loss differences favor the augmented model.

| Metabolite | In-sample $\Delta L$ | LOTO $\Delta L$ | Unadjusted $P$ | FWER $P$ |
|---|---|---|---|---|
| Glycocholate | 0.821 | 0.497 | 0.059 | 0.212 |
| Lactate | 0.689 | -0.443 | 0.038 | 0.294 |
| Propionate | 0.294 | -0.704 | 0.032 | 0.639 |
| Indole-3-propionate | 0.276 | -1.873 | 0.179 | 0.678 |
| Imidazolelactate | 0.173 | -1.750 | 0.352 | 0.853 |
| Succinate | 0.152 | -0.017 | 0.260 | 0.865 |
| Cholate | 0.049 | -2.133 | 0.585 | 0.968 |
| Lithocholate | 0.019 | -1.001 | 0.599 | 0.995 |

The FWER probability compares each observed loss improvement with the maximum improvement across all eight metabolites under each of 2,401 within-subject circular-shift combinations. Influence analysis further showed that omitting H4001 changed the glycocholate coefficient from negative to positive and its loss improvement from 0.821 to -0.075. Lactate behaved similarly, with loss improvement changing from 0.689 to -0.042. Thus, the pooled coefficients are driven partly by a single subject. A hierarchical random-effects model was not used because four clusters cannot support reliable estimation of a population random-effect distribution.

In the secondary 10-subject cohort defined by at least six usable transitions, glycocholate and lactate remained the first and second in-sample candidates. This ranking agreement is descriptive and is not used as confirmatory evidence because the expanded cohort has heterogeneous transition counts and was introduced as a sensitivity analysis.

### S8 Antibiotic and other perturbation metadata

Categorical metadata were encoded from their Yes/No values before numerical analysis. Two of 32 retained observations were antibiotic-positive: one in C3032 and one in C3037.

The sensitivity leaves only five transitions for each exposed subject and is not suitable for subgroup inference. Diarrhea was recorded in multiple retained observations, and one hospitalization was recorded. We therefore avoid the broad statement that all major perturbations were rare. The defensible limitation is that the number of subjects and transitions is insufficient to separate these time-varying exposures from underlying dynamics.

**Table S9:** Sensitivity excluding transitions adjacent to antibiotic-positive samples.

| Subject | Positive observations | Excluded transitions | $b$ all | $b$ after exclusion |
|---|---|---|---|---|
| C3032 | 1 | 2 | -1.356 | -1.663 |
| C3037 | 1 | 2 | 0.536 | -0.222 |
| H4001 | 0 | 0 | -1.034 | -1.034 |
| H4004 | 0 | 0 | -0.794 | -0.794 |
